# Modeling epistasis in mice and yeast using the proportion of two or more distinct genetic backgrounds: evidence for “polygenic epistasis”

**DOI:** 10.1101/555383

**Authors:** Christoph D. Rau, Natalia M. Gonzales, Joshua S. Bloom, Danny Park, Julien Ayroles, Abraham A. Palmer, Aldons J. Lusis, Noah Zaitlen

## Abstract

**Background:** The majority of quantitative genetic models used to map complex traits assume that alleles have similar effects across all individuals. Significant evidence suggests, however, that epistatic interactions modulate the impact of many alleles. Nevertheless, identifying epistatic interactions remains computationally and statistically challenging. In this work, we address some of these challenges by developing a statistical test for *polygenic epistasis* that determines whether the effect of an allele is altered by the global genetic ancestry proportion from distinct progenitors.

**Results:** We applied our method to data from mice and yeast. For the mice, we observed 49 significant genotype-by-ancestry interaction associations across 14 phenotypes as well as over 1,400 Bonferroni-corrected genotype-by-ancestry interaction associations for mouse gene expression data. For the yeast, we observed 92 significant genotype-by-ancestry interactions across 38 phenotypes. Given this evidence of epistasis, we test for and observe evidence of rapid selection pressure on ancestry specific polymorphisms within one of the cohorts, consistent with epistatic selection.

**Conclusions:** Unlike our prior work in human populations, we observe widespread evidence of ancestry-modified SNP effects, perhaps reflecting the greater divergence present in crosses using mice and yeast.

**Author Summary:** Many statistical tests which link genetic markers in the genome to differences in traits rely on the assumption that the same polymorphism will have identical effects in different individuals. However, there is substantial evidence indicating that this is not the case. Epistasis is the phenomenon in which multiple polymorphisms interact with one another to amplify or negate each other’s effects on a trait. We hypothesized that individual SNP effects could be changed in a polygenic manner, such that the proportion of as genetic ancestry, rather than specific markers, might be used to capture epistatic interactions. Motivated by this possibility, we develop a new statistical test that allowed us to examine the genome to identify polymorphisms which have different effects depending on the ancestral makeup of each individual. We use our test in two different populations of inbred mice and a yeast panel and demonstrate that these sorts of variable effect polymorphisms exist in 14 different physical traits in mice and 38 phenotypes in yeast as well as in murine gene expression. We use the term “polygenic epistasis” to distinguish these interactions from the more conventional two- or multi-locus interactions.

## Introduction

Genetic association studies in humans and model organisms have identified a number of significant links between individual polymorphisms and phenotypic variability. A fundamental assumption of many of these studies is that an allele will have a similar effect in each member of the population, that is, that epistatic and other higher-order interactions across the genome can largely be ignored(1–4). Prior studies to specifically detect epistasis in flies(5), yeast(6), mice(7,8), and humans(9) have considered pairwise interactions between loci to begin to identify interacting loci, yet this approach is hampered by a substantial increase in the threshold of significance due to the increased numbers of tests (from n to at least 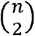, where n are the number of SNPs tested). In this study, we consider detection of epistasis by testing the effects of interactions between individual polymorphisms and global ancestry (θ), which we defined as the percentage of the genome inherited from a given ancestor or ancestral population.

Recently, we developed a SNP-ancestry interaction model and applied it to a human dataset, detecting only modest evidence of ancestry-specific genetic effects(10). There is substantial evidence of interactions with ancestry in model organisms. For example, we observed radically different phenotypic consequences of null alleles of *Tcf7l2* and *Cacna1c* when expressed on different inbred strain backgrounds(11). Recently, a number of powerful genetic resources have been developed for model systems to map variation in complex traits, such as the Hybrid Mouse Diversity Panel(12), the Collaborative Cross(13,14), the BXD recombinant inbred panel(15), various outbred mouse and rat populations(16), or the Drosophila Synthetic Population Resource(17). The variation in ancestry present in these populations provide a unique opportunity to test for ancestry-driven epistasis, but no existing method is suitable for this task.

To address this problem, we incorporate a linear mixed model capable of accounting for the highly structured relatedness of constructed model organism(18) into our previous approach. We then examine two distinct mouse cohorts as well as a panel of yeast crosses and test for interactions between the effects of individual polymorphisms and the percentage of the genome for each individual that originated with a specific founder. Although our initial model was designed to identify both genetic background and environmental effects, the controlled environment in which model organisms are raised is intended to minimize sources of environmental variance. This test, which we have called Gxθ, acts as a surrogate measure of the concurrent action of many other SNPs, and allows us to ask whether or not the effect size of a given SNP changes as a function of overall genetic ancestry. In contrast to a pairwise epistatic test, where significance means a detected interaction between two specific loci, a natural interpretation of a significant Gxθ interaction is widespread epistasis of the tested genotype G with many loci across the genome, as we previously showed(10).

We first evaluated the statistical properties of our method on simulated phenotypes based on real recombinant inbred line data sets(12). The Gxθ test successfully distinguishes between the marginal effects driven only by a given SNP or ancestry and effects that arise from their interaction. We then applied the test to two different populations of mice: recombinant inbred (RI) lines that are a subset of the HMDP(12) (143 phenotypes), and a 50^th^ generation intercross between the inbred LG/J and SM/J mouse strains(19), whose ancestry proportions have been clearly and precisely determined(19) (75 phenotypes). To show that our approach is broadly applicable, we also apply this test to a yeast panel consisting of thousands of progeny derived from 15 distinct yeast crosses(20) (40 phenotypes). Utilizing gene expression data from the HMDP cohort in two different conditions, we further demonstrate that our approach replicates findings across similar populations.

Given this evidence of epistasis, we examined the ancestry distribution at specific sites in the mouse HMDP RI lines. Under neutrality, on average, we would expect equal representation of founder ancestry in the panel, however, we observed that for many positions in the genome there is a statistically significant depletion or overrepresentation of specific founder ancestries. We interpret this as evidence of selection during the process of RI strain derivation and we identify regions of the genome with strong selection up to 4 Mb in length. These regions are evidence of strong Gxθ interactions which inhibit or promote the transmission of specific alleles to subsequent generations. These regions are enriched for genes involved in cancer and organogenesis and are enriched (P=1.8E-4) for metallopeptidases, which play key roles in fertility and neo/perinatal lethality(21–23).

Our observation of genomic locations where the effect size of a given polymorphism is driven by overall genomic context and ancestry highlights the importance of studying epistasis and other effects in model organisms. Studying these context-dependent effects in human cohorts has been notoriously difficult, but our results indicate that given the right design and analytical framework these effects can be highly significant, and has implications for our ability to build predictive models from genotype to phenotype and account for ancestry effects in population studies.

## Results

### Method Overview

Our objective is to determine whether a given SNP i has a different effect size as a function of changes in overall genetic background. As described in our previous work(10), this is effectively achieved through an interaction test between each SNP and global genetic ancestry θ. In a standard test of the association between a polymorphism ***X_i_*** and a phenotype **y**, we can model this relationship as ***y*** = *μ* + *β_i_**X**_**i**_* + ***u*** + ***e***, where μ is the mean phenotypic value, β is the effect size of the SNP, u is a random variable accounting for relatedness and e is a random variable accounting for all other sources of error in the study. We extend this analysis to account for interactions between a polymorphism and ancestry by adding terms to our model such that the relationship now becomes

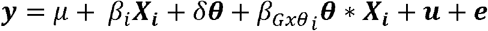

where δ is the global weight of the ancestry effect, θ are the ancestries for all N individuals and *β_Gxθ_* are the weights of the Gxθ effect. Our Gxθ test is then an LRT test with a null of *β_Gxθi_* = 0 and an alternate of *β_Gxθi_* ≠ 0.

To account for multiple testing, we follow the best practices of the respective model organism communities, specifically we use the previously reported genome-wide significance threshold for the HMDP of 4.2E-6 (24) or the AIL of 8.06E-5 (19). For Yeast, we follow the example previously reported of Bloom et al (25) and use an FDR of 5%.

### Simulated Data

To examine the properties of our approach in a model organism cross, we applied the method to phenotypes simulated using real genotypes that reflect the underlying relatedness present within the recombinant inbred (RI) lines of the Hybrid Mouse Diversity Panel (HMDP). Random SNPs with a minor allele frequency (MAF) between 25% and 75% were selected from our panel and phenotypes constructed at 200 values of either a main SNP effect β_G_ or a SNP-ancestry interaction effect β_Gxθ_ where θ is the proportion of all SNPs arising from C57BL/6J (see Methods) mice for interaction with each individual SNP with 1,000 simulations at each β (200,000 total simulations) (see Methods).

We observe that changes to either β_G_ or β_Gxθ_ independently did not affect the power of identifying an association for β_Gxθ_ or β_G_, respectively, indicating that these two terms are correctly being estimated independently of one another. We briefly explored incorporating a second genetic relationship matrix (GRM) accounting for population structure arising from descent from a single ancestor (the B6 mouse)(26), however we observed no significant improvement in power when incorporating this second GRM (Figure 1a, 1b).

**Figure 1.**
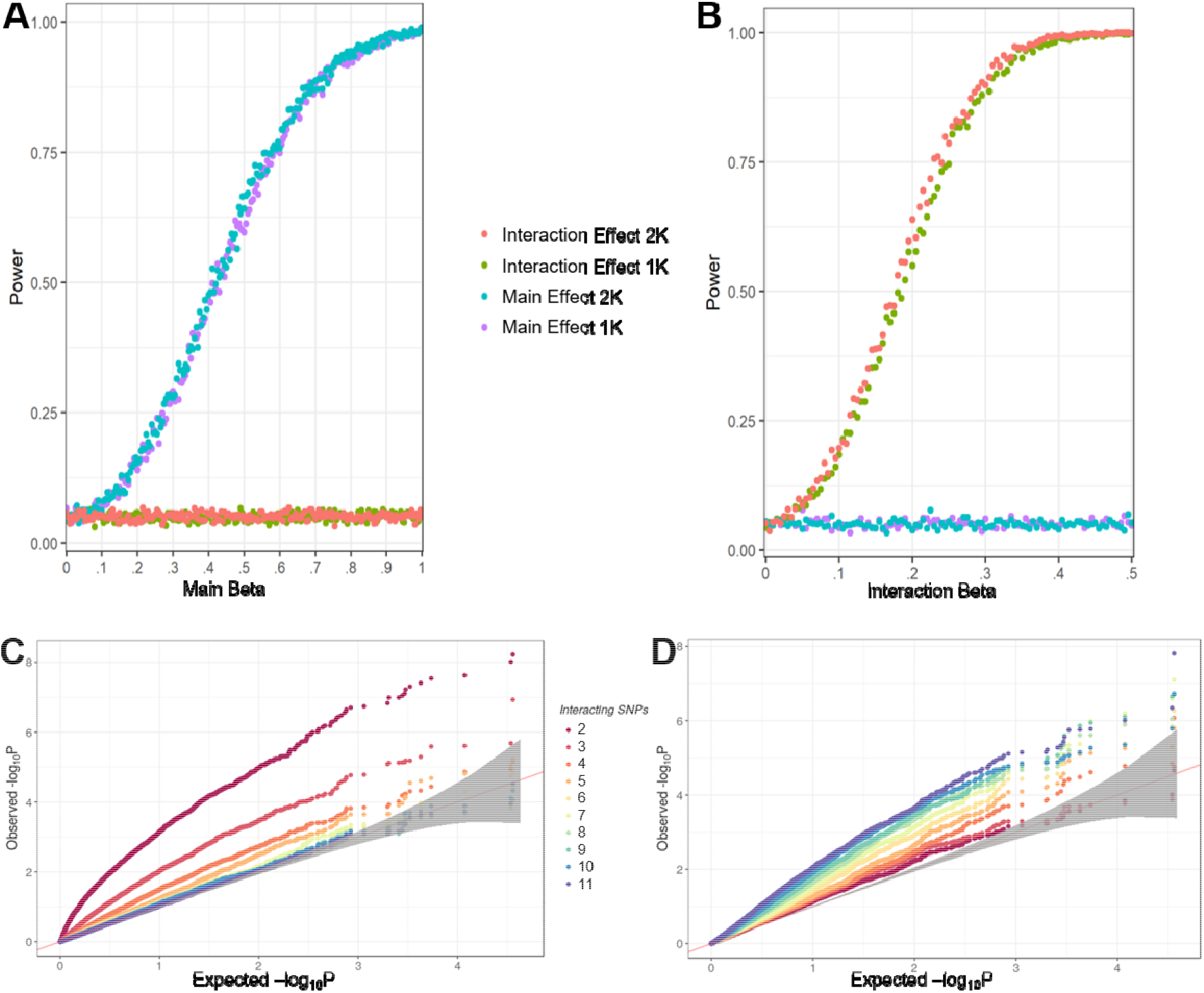
Simulation Results. A+B) Power calculations based on simulated data with variable main SNP effects β_G_ (A) or variable Ancestry-SNP effects β_Gxθ_ (B). Blue and Purple power curves are the power curves for detecting a significant SNP effect, while the orange and green curves are the power curves for detecting a significant Gxθ effect. Two phenotypic models, one incorporating 1 GRM (1K) correcting for relatedness in the SNPs (green, purple) and one incorporating 2 GRMs (2K) correcting for relatedness in both SNPs and Ancestry (red, blue) were used. (C+D) Effect of increasing numbers of epistatically interacting SNPs on the algorithms ability to detect a significant main SNP effect (C) or Gxθ effect (D). The straight line in C and D represents the line where expected and observed P values are the same, while the grey band indicates the region in which the P-value distribution cannot be distinguished from the expected P-value distribution.

To evaluate what forms of interactions the test is powered to identify we examined a range of epistatic architectures, focusing specifically on our method’s ability to recover higher-order interactions, a type of epistasis which traditional models struggle to accommodate. For 100,000 simulations, a SNP was selected at random from SNPs with MAF between 25 and 75% and combined with 1-10 additional SNPs (10,000 each) and phenotypes generated for these SNP groupings followed by analysis by the Gxθ algorithm (see Methods). As the number of interacting SNPs increases, the power to detect the individual SNP decreases as expected (Figure 1c), however the power to detect a Gxθ locus rises (Figure 1d), suggesting that our model is powerfully situated to identify SNPs involved in complex higher-order interactions, especially those consisting of more than 5 interacting partners (Figure S1).

### Application to *in vivo* Data

#### Mouse Populations

We applied the Gxθ test to two large panels of mice to identify Gxθ effects (i.e. instances where a given polymorphism interacts epistatically with one or many other loci as captured in the model by θ, the global ancestry). Our first cohort is the HMDP, a set of 150+ commercially available inbred strains(27). Numerous GWAS have been performed in the HMDP, including several using PYLMM, which forms the core of our algorithm(24,28,29). The largest component of the HMDP is comprised of 122 RI strains (28 AxB, 71 BxD, 12 BxH, 11 CxB). Each RI was constructed from re-derivation of novel inbred lines via brother-sister mating following an F2 cross between the sub-panel’s parental lines. In the case of the HMDP, one of the parental strains for each RI strain was the commonly studied strain C57BL/6J (**B6**). Using B6 as the common ancestral line to the RI panels, we calculated θs for each RI strain (Fig 2a). As expected, the average ancestry attributable to B6 was roughly 50% (50.63%) and roughly normally distributed. We removed a single outlier, BXD32/TyJ, whose B6 Ancestry of 25.41% reflects a previously known additional backcross to DBA/2J, resulting in a strain that is 75% DBA and 25% B6. Each study using the HMDP uses a different subset of the entire panel, and we selected a study on heart failure induced by the chronic beta adrenergic agonist isoproterenol(30) for analysis as it used the most RI strains compared to other published HMDP data. We used 123 clinical phenotypes in conjunction with microarray-derived gene expressions measured in the left ventricle and on average tested 67 RI strains per phenotype.

**Figure 2.**
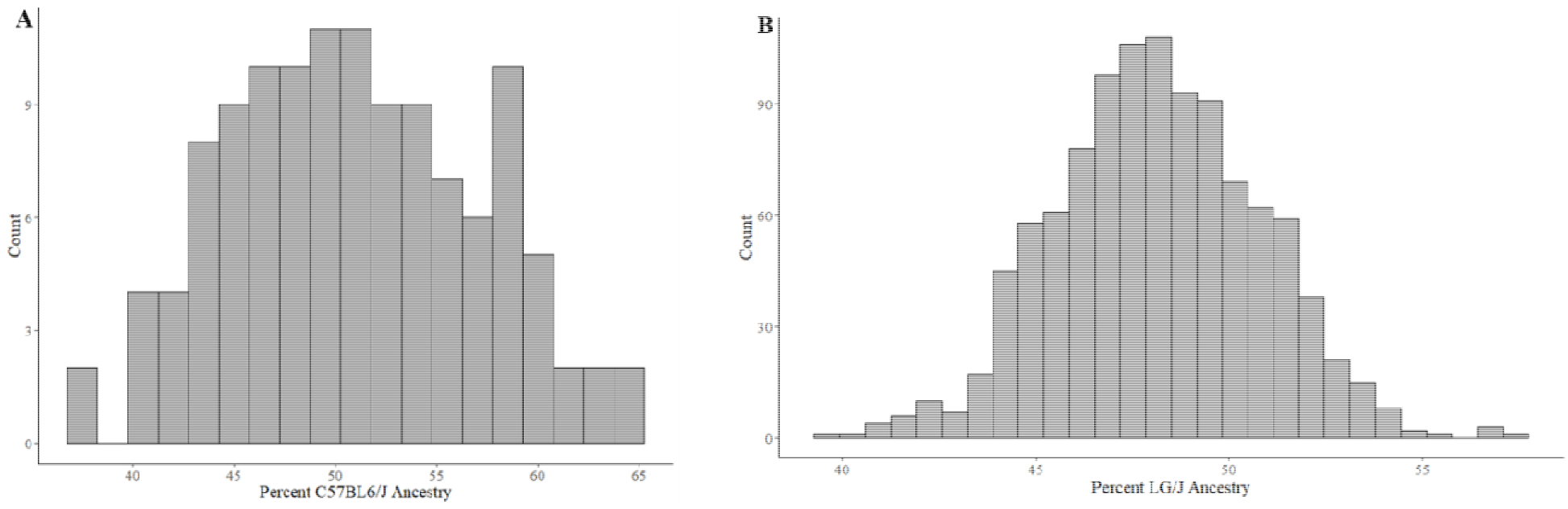
Ancestral Strain Contributions by Strain. **A)** The 122 strains of the RI panel of the HMDP, **B)** The 1063 animals in the F_50_ – F_56_ generation of the AIL Cross

The second cohort consists of 1,063 animals from the F_50_ – F_56_ generation of an advanced intercross line (**AIL**) created by crossing the LG/J and SM/J inbred mouse strains (19). Unlike the RI strains from the HMDP, AILs are maintained in a manner that minimizes inbreeding and promotes genetic diversity at each generation. We arbitrarily set LG/J as the ancestral strain of interest and calculated θs for each of the 1,063 mice in the panel (Fig 2b). For this study, we focused on a diverse set of 133 phenotypes that had been measured in these mice. We describe the results of the Gxθ associations in each panel before demonstrating replication of interactions in a phenotypic trait as well as in expression data.

#### Evidence of SNP x Ancestry Interactions in Phenotypic Data

We applied the Gxθ method to the 123 observed heart-failure related phenotypes from the HMDP RI strains. We observed well-calibrated statistics, with λ_GC_ equal to 0.978 and λ_GxθC_ equal to 1.045 after pooling the p values across all phenotypes. We observed 44 significant Gxθ loci across 9 phenotypes: E/A ratio, free fatty acid content in the blood, cardiac fibrosis, fractional shortening of the heart during contraction, heart rate, internal diameter of the left ventricle, left ventricular mass and left and right atrial weights (Table 1 and S1). These Gxθ loci were largely distinct from previously reported GWAS loci in the same phenotypes in this panel of mice(28,31), yet contained a number of highly relevant genes, as discussed below.

**Table 1.** Most significant Gxθ associations for each phenotype observed in the AIL or HMDP cohorts as well as all Yeast Gxθ loci passing a cross-wide 5% FDR threshold See table S1 for complete list of loci. Act3.2 is ‘activity levels in control animal on day 3 of a conditioned place preference test’, LVIDs is ‘left ventricular internal dimension at systole’ and PT is post isoproterenol treatment.

| Cohort | Phenotype | Chromosome | Basepair | rsID | P |
| --- | --- | --- | --- | --- | --- |
| AIL | Act3.2 | 11 | 93607101 | rs27076868 | 6.22E-07 |
| AIL | Average Body Weight | 6 | 133867165 | rs30057768 | 1.62E-06 |
| AIL | Glucose | 9 | 70318603 | rs50320206 | 3.03E-07 |
| AIL | Weight at 68 days | 6 | 133899245 | rs47230944 | 2.09E-06 |
| HMDP | E wave 3 weeks PT | 19 | 21360156 | rs36743940 | 5.71E-10 |
| HMDP | E/A ratio 3 weeks PT | 6 | 8462274 | rs47261338 | 7.10E-08 |
| HMDP | Fibrosis control | 17 | 41256115 | rs6295287 | 6.74E-07 |
| HMDP | Fractional shortening control | X | 48286693 | rs30272504 | 3.69E-06 |
| HMDP | Free fatty acids control | 11 | 113035483 | rs27020574 | 4.09E-06 |
| HMDP | Heart rate 1 weeks PT | 18 | 9408717 | rs29769121 | 2.37E-06 |
| HMDP | Heart rate control | X | 48286693 | rs30272504 | 2.68E-06 |
| HMDP | Left atrial weight 3 weeks PT | 16 | 54550810 | rs51319671 | 2.18E-06 |
| HMDP | LVIDs control | X | 48286693 | rs30272504 | 2.92E-06 |
| HMDP | Left ventricular mass 1 weeks PT | 3 | 148140470 | rs31313229 | 2.91E-06 |
| HMDP | Left ventricular mass 2 weeks PT | 17 | 72564961 | rs50549031 | 8.65E-08 |
| HMDP | Right atrial weight 3 weeks PT | 16 | 49837267 | rs50479702 | 4.13E-07 |
| Yeast | Cross 2999: EGTA | XIV | 622639 | NA | 2.91E-07 |
| Yeast | Cross 2999: Lithium Chloride | IV | 519675 | NA | 1.04E-07 |
| Yeast | Cross 2999: Lithium Chloride | XII | 341678 | NA | 1.16E-05 |
| Yeast | Cross 2999: Manganese Sulfate | XIV | 622639 | NA | 2.30E-06 |
| Yeast | Cross 3043: Manganese Sulfate | XVI | 206053 | NA | 4.17E-10 |
| Yeast | Cross 3043: Paraquat | XV | 410166 | NA | 3.14E-08 |
| Yeast | Cross 3043: YNB | XV | 561457 | NA | 3.67E-07 |
| Yeast | Cross 3043: YPD | XV | 561457 | NA | 2.50E-06 |
| Yeast | Cross B: Maltose | VII | 1067754 | NA | 1.20E-07 |

By way of example, we focus on two important phenotypes from the HMDP panel. Cardiac fibrosis is a marker of cardiac dysfunction. Genes identified through the Gxθ screen (Figure 3B) as potential candidates for cardiac fibrosis include: *Crisp2* (rs6295287, p=6.74E-7), a secreted biomarker of cardiovascular disease(32), *Top2b* (rs31538570, p=2.14E-06), which plays a cardioprotective role in response to stress(33), *Rarb* (rs31538570, p=2.14E-06), a known regulator of inflammation with unknown function in the heart(34), and *Fibrosin* (rs33146511, p=2.54E-6), a major component of the fibrosis pathway(35). An increase in left ventricular mass in response to catecholamine challenge is the primary marker of cardiac hypertrophy in the HMDP. A single Gxθ locus(rs31313229, p=2.91E-06) on chromosome 3 (Figure 3C) contains the gene *Lphn2*, which has a role in the promotion of cellular adhesion in response to external stimuli.(36) Crucially, none of these loci, many of which contain candidates of particular interest or relevance to heart failure, were reported in the original study, suggesting the possibility of new targets for further therapeutic research.

**Figure 3.**
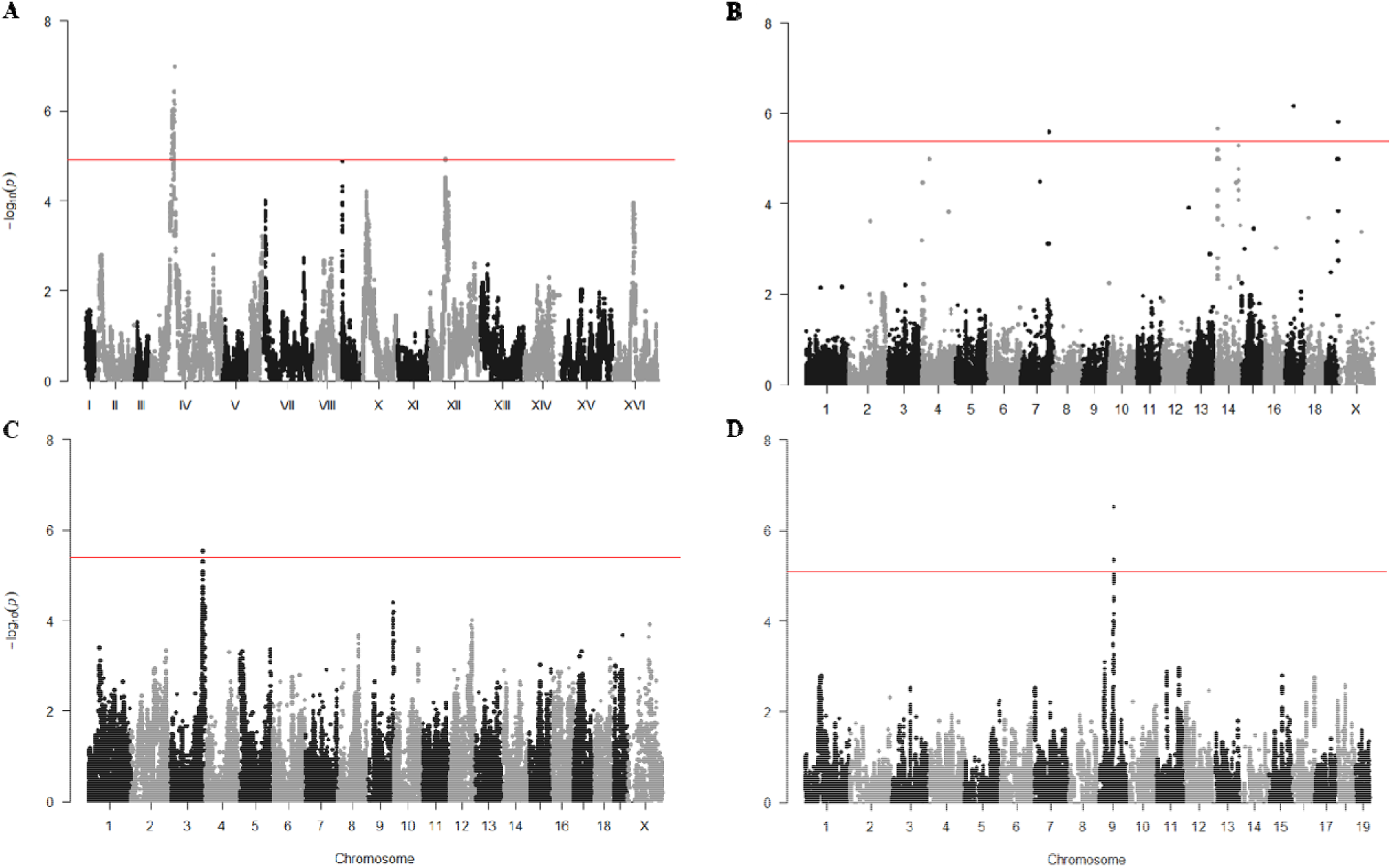
Gxθ Results in the Yeast, HMDP and LGxSM AIL Cohorts. A) Lithium Chloride Exposure in Yeast B) Cardiac Fibrosis and C) Left Ventricular Weight in the HMDP D) Glucose in the AIL. Significance Threshold in Yeast (A)= 1.2E-5 (FDR 5%) HMDP (B,C)= 4.2E-6(24), AIL (D)=9.1E-6 (FDR 5%)

We next applied the Gxθ method to 133 phenotypes measured in the LG/J x SM/J AIL, including the sex of the animal as a covariate as, unlike the HMDP data, the AIL consists of both male and female animals. As with the HMDP data, we observed well-calibrated statistics in this cross, with λ_GC_ equal to 0.995 and λ_GxθC_ equal to 1.033. Despite the larger sample size, we observed only 4 significant loci across 4 phenotypes: activity levels in a saline-injected animal on day 3 of a conditioned place preference test; average weight of animals across 5 different time points roughly a week apart; glucose (mg/dL) in blood after a 4 hour fast [Figure 3D]; weight at ~68 days of age (Table 1 and S1).

#### Yeast Populations

We also used the Gxθ algorithm to examine 15 previously reported yeast crosses(20) ranging from 650 to 950 progeny per cross for evidence of significant Gxθ loci and an average of 71,000 segregating genetic variants. We examined the responses of these progeny to forty different chemical stimuli(20) and recovered 92 loci across 38 phenotypes which met a 5% FDR threshold for that cross/phenotype combination (Table S2). At a more stringent level of significance (5% FDR across an entire cross), we identify 9 loci across 7 phenotypes (LiCl, EGTA, Maltose, MnSO4, Paraquat, YNB, YPD) which show strong evidence of Gxθ interactions (Table 1, Figure 3A).

### Gxθ Associations in HMDP Cohort Gene Expression

We next examined gene expression in the hearts of the HMDP cohort using previously obtained transcriptome microarrays from both control and treated conditions(37). Each cohort consists of approximately 70 RI lines, with 66 lines overlapping between the two cohorts (full lists of strains in Table S2). We examined 13,155 expressed and varying (Coefficient of Variation > 5%) genes from the left ventricles of the HMDP cohorts using the ~170k SNPs with MAF >= 0.05. We observed 1,486 significant associations with 18 genes at a Bonferroni-corrected P value of 3.2E-10 (135,130 associations with 1,350 genes at GW-significant threshold of 4.2E-6) in the control cohort and 597 significant associations with 39 genes at the same threshold in the treated cohort (32,043 associations with 1,042 genes at the genome-wide significant threshold of 4.2E-6(24)) (Full results available in Table S10).

Phenotypic variability was larger in the catecholamine treated cohort of the heart failure HMDP study (average Coefficient of Variation in control was 22.6% vs 28.6% for the treated cohort)(30), which could partially explain the lower number of significant associations observed in that group of mice. Notably, we saw no enrichment for SNPs in the *cis*-regions of the examined genes in contrast to what is observed in standard mapping. Genes in the control cohort with significant Gx□ associations are enriched for mitochondria-associated genes (P=1.8E-10 (P=.032 in treated)), suggesting a role of Gx□ interactions in the regulation of mitochondrial dynamics. Genes in the treated cohort with significant associations were enriched for genes involved in post-transcriptional modifications to RNA including RNA splicing (P=7.4E-3), highlighting the importance of alternative splicing to the response to catecholamine challenge(38).

#### Replication of Gx□θ Associations across eQTLs from HMDP Cohort

To demonstrate that the method is able to replicate Gxθ results across cohorts, we examined the reproducibility of expression Gxθ QTLs in the treated and untreated RI lines of the HMDP. Of the 1,486 associations observed in the control data, we observe 305 (21%) with Gxθ interaction (P<.05) in the treated cohort (36 at FDR< 5%), indicating a strong replication (p = 2.2E-16 from a binomial test) between the two cohorts despite differences in genetic background and the potentially reduced power in the treated cohort caused by the effects of the catecholamine drug.

### Distortion from Expected Allelic Frequencies

After population admixture, allelic frequencies will shift in the resulting population. This can be due to the effects of selection against deleterious allelic combinations (i.e. fitness epistasis), meiotic drive (i.e. intragenomic conflict resulting in the transmission of one or more alleles over another during meiosis). Alternatively, allele frequencies may shift through random genetic drift. We hypothesize that Gxθ loci can be driven by fitness epistasis, or instances where allelic variation in one locus affects fitness at other loci across the genome. This hypothesis is in line with prior research(39–41) which has explored allelic imbalance or gametic ratio distortion as a function of epistasis and fitness. Examining the genomes of admixed population for signs of significant allelic shift may reveal interesting loci that can inform long standing questions in evolutionary and population genetics about the forces that maintains variation in fitness in populations, as well as the presence of non-equal chromosomal inheritance.

We therefore conducted a test to search for individual loci with enriched or depleted B6 ancestry in the RI strains. As expected, the average B6 ancestry across all SNPs was 50.68% +/− 8.1%, which is indistinguishable from the ancestry by strain average of 50.63% +/− 6.7% (Fig 4a). At the level of individual loci, however, we observed significant variation in B6 ancestry across the genome (Fig 4b), with some SNPs displaying very low or very high frequency (Fig 4c). We calculated the statistical likelihood of detecting the observed ancestries at all loci across the genome (λ=1.07, Fig 4d).

**Figure 4.**
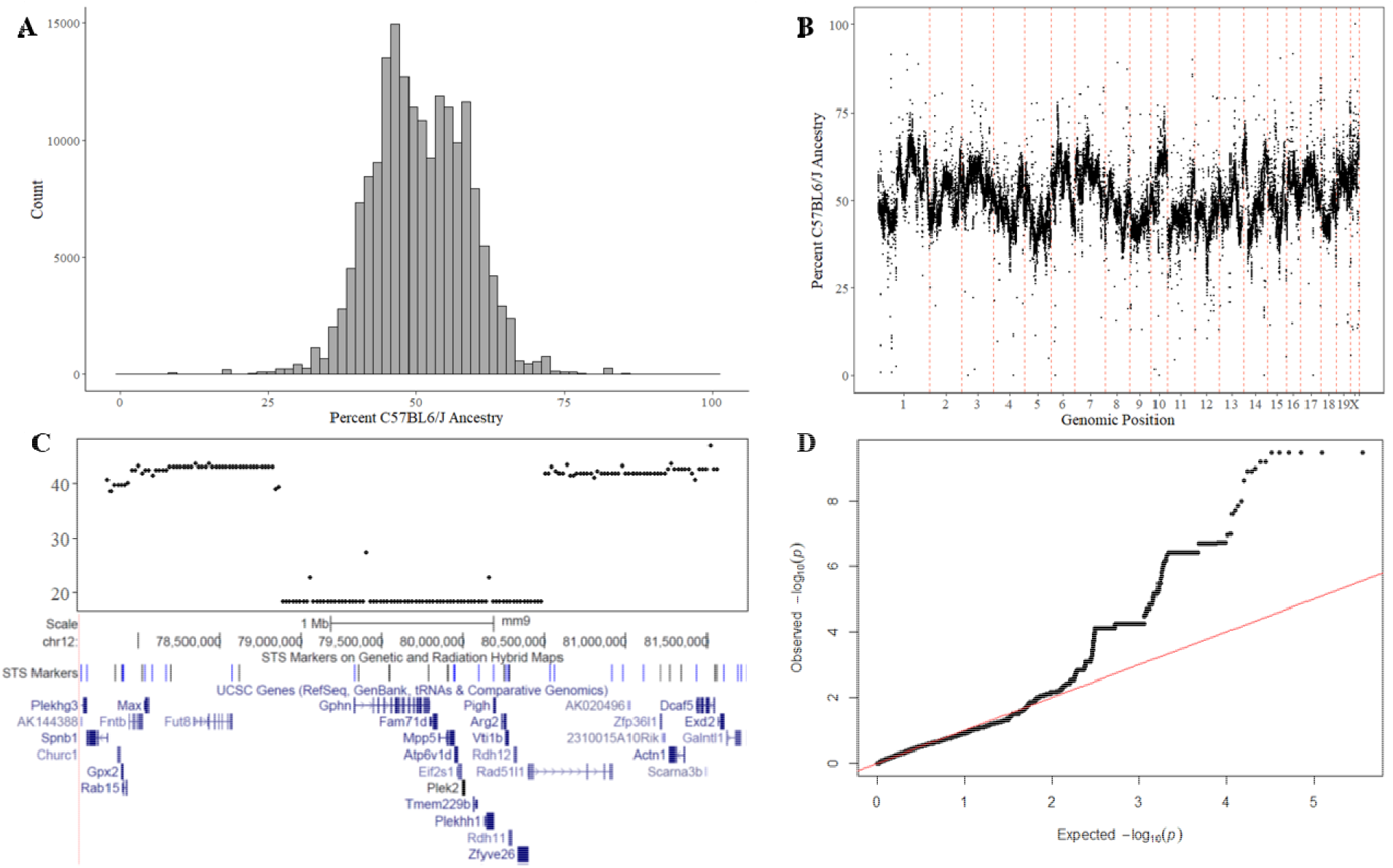
Allelic Frequency Imbalances of individual sites across the genome. **A)** A histogram of B6 ancestry across the geneome. **B)** Genome-wide plot of ancestry **C)** Zoomed in region on chr 12 with significant depletion of B6 allele. **D)** QQ plot of all Pvalues. Genomic Inflation Factor λ=1.07

We subsequently performed a Wald test for ancestry frequency (θ) at each SNP i where 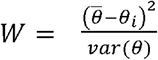. We used the global variance in the denominator, which assumes that most SNPs lie under the null and which is conservative if this assumption is violated and could be replaced with a more powerful test. For each SNP, we examined strains which were not invariant for that locus and identified 614 SNPs from the 170k original SNPs with significantly altered ancestries at an FDR of 5% (Table S3). We suspect that these SNPs do not represent passive effects such as genetic drift due to the nature of how RI panels are created, with a single F2 generation of mice followed by brother-sister mating and eventual fixation as similar effects on allelic balance have been attributed to genetic incompatibilities in other mouse populations(42). Allelic balance of paternal alleles across the RI panel, therefore, may be modeled using a Bernoulli distribution. In the HMDP, with 122 RI lines, the odds of a 30% imbalance (80% B6 vs 20% other) would be 8.3E-12, which, even across 200,000 SNPs results in 2E-6 expected SNPs with this level of imbalance across the entire genome.

Nine loci were identified in which 5 or more SNPs were located together (Table 2). These 9 regions contain the majority (525, 86%) of all significant affected loci and indicate regions where significant allelic imbalances are occurring. One region of particular interest lies on chromosome 18 and contains 246 SNPs (40% of all significantly altered SNPs) with an average B6 ancestry of 82.5%. SNPs in this region are found between B6 and C3H/HeJ and B6 and BALB/cJ. This locus contains a number of interesting candidate genes potentially relating to organismal fitness, including *Epc1*, a transcription factor linked to DNA repair, muscle differentiation and cancer suppression(43,44). It is also the only gene within the locus whose expression has a suggestive Gxθ association with the locus itself (P=0.014).

**Table 2.**
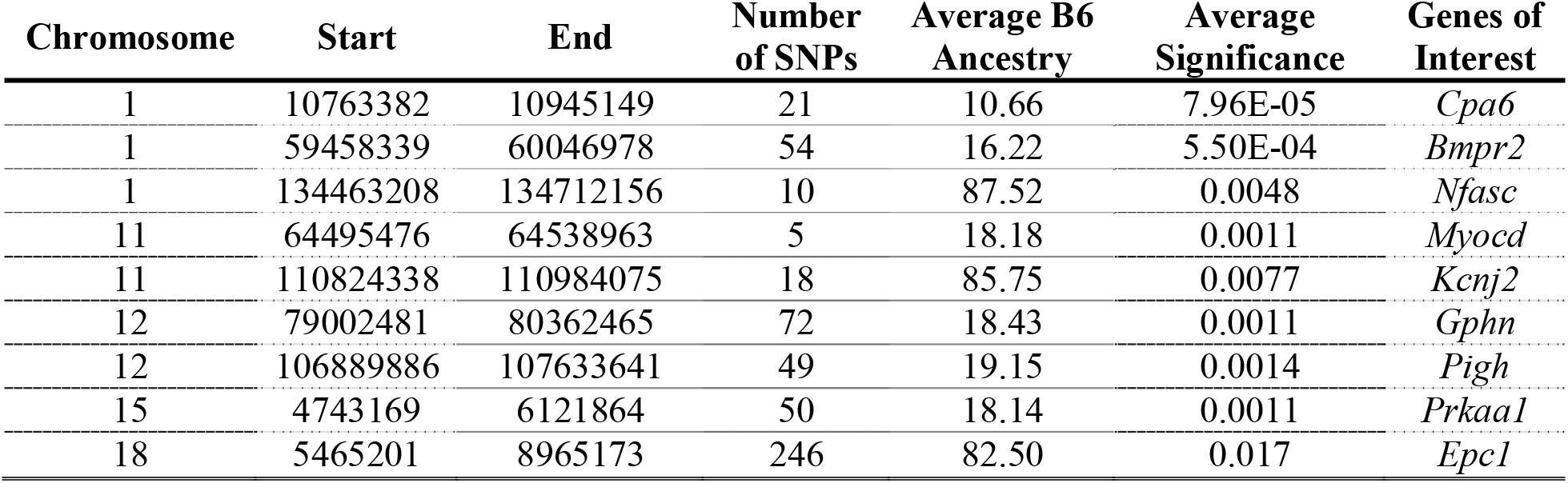
Regions with Significant and Sustained B6 Depletion or Selection

When all genes within 2 MB (the average Linkage Disequilibrium (LD) block size in the HMDP(27)) of boundaries of these allelic imbalance loci are examined as a whole, we observe significant enrichments for genes involved in cancer (corrected P-values range from 1.3E-8 to 1.6E-7). These genes, whose functions range from cell cycle control to organismal growth and DNA damage mediation and repair are not unexpected given their crucial role in organismal survival. If the B6 or alternative allele deleteriously affect the function of one or more of these genes when combined with new alleles from other strains located elsewhere on the genome, then it is reasonable to assume that the ability for an individual to pass on its genetic information to the next generation may be significantly reduced. Other enriched gene ontology categories include the crucial-for-life categories of respiratory tube development (2.7E-4) and heart development (7.6E-4), likely mediated through changes in Matrix Metalloproteinase activity (3.9E-8). These enzymes are canonically responsible for the regulation of the extracellular matrix, but have been linked to the modulation of responses to many bioactive molecules and have important roles in fertility, embryonic development and neo/perinatal mortality in addition to many other diseases(23). Taken together, these enrichments suggest that we may be observing depletion due to epistatic interactions with other loci which were neutral or beneficial in the original inbred lines, but which become deleterious for a given allele when placed into a new genetic environment.

## Discussion

We present a test which we call Gxθ, that leverages admixed populations such as inbred mouse strains to identify sites of “polygenic epistasis” where a SNP interacts with an unknown number of other loci summarized into a single genomic ancestry score, θ. One major advantage of this approach is that, unlike other epistasis-focused association approaches, it does not increase the number of tests when compared to a typical GWAS, so similar significance thresholds may be used.

The existence of epistatic interactions and their role in human diseases and phenotypes have been known for many years, with notable examples of gene-gene interactions in Alzheimer’s disease(45), Bardet-Biedl syndrome(46) as well as classic interactions governing hair color and skin pigmentation. Despite these examples and many more like them, epistatic interactions have proven difficult to find using association techniques(47). One notable exception has been the success of using consomic strains of mice, where one or more chromosome in strain A is replaced with a chromosome from another strain B(48). This approach, which increases or decreases the contribution of a given strain to phenotypic and genetic variance in a clear and controlled manner has resulted in the identification of numerous epistatic interactions(49,50).

Our method sought to circumvent this traditional pitfall in Epistasis GWAS by examining each SNP only once for an interaction with a global genomic ancestry to identify interactions or “polygenic epistasis”. Our model is able to identify much higher order interactions than can be observed using traditional SNP-SNP interaction models. As demonstrated through simulation, our model in fact increases in power as the complexity of the epistatic interactions increases, likely reaching peak power in the hypothetical scenario where every SNP interacts with the tested SNP to affect phenotypic expression. When applied to human populations, a different form of our test was able to find several significant associations(10)

In this study, we examined two populations of mice: the recombinant inbred lines of the Hybrid Mouse Diversity Panel(27) and an Advanced Intercross Lines (AIL) based on LG/J and SM/J(19) as well as a yeast RIL mapping panel founded by fifteen yeast strains(20). We observed significant Gxθ interactions in 14 of the 15 yeast crosses that affected over half of the examined phenotypes, reinforcing the widespread role of genotype-by-genotype interactions in regulating quantitative traits across diverse genetic backgrounds and phenotypes. In the mouse cohorts, however, despite nearly ten times the number of genetically distinct mice as well as a larger number of phenotypes and genotypes in the AIL, we observed approximately eleven-fold more significant Gxθ peaks in the HMDP phenotypes than in the AIL phenotypes (44 vs 4).

Several reasons may account for this difference. First, outbred mice have significantly lower power due to increased phenotypic variance caused by high rates of heterozygosity at alleles while inbred populations have increased power due to a lack of this heterozygosity. This can also be seen in our regular GWAS results(19,30,31). Second, the different phenotypes studied in each cohort will have different genetic architecture and experimental noise (e.g. the AIL study includes many behavioral traits while the HMDP does not). Third, significantly more recombination events have occurred in the AIL compared to the HMDP, leading to a tighter distribution of background and a larger number of independent tests. Fourth, the genetic backgrounds of the two cohorts are different, which might also contribute to differences in observed numbers of Gxθ interactions. Although individually, LG and SM are only slightly less genetically diverged from one another as any pair of strains that make up the RI panels of the HMDP (Table S4), the presence of five ancestral lines in the HMDP compared to only two in the F50 cross results in much more genetic diversity in the HMDP compared to the F50 mice. Finally, we observe a higher variance in ancestral background in the mice HMDP when compared to the AIL. As our method relies on differences in ancestral background to identify sites with different effect sizes in different genetic contexts, the reduced variance in the AIL lines necessarily corresponds to a decrease in the power to detect Gxθ interactions. Taken together, our method is best suited to datasets with relatively low heterozygosity, clear and numerous differences in genetic background, and with higher variance in the percentage of SNPs attributable to a given ancestry.

Furthermore, we identify nine regions across the genome of the RI lines of the HMDP which show a strong selection for or against a particular parental allele. We propose that this is a result of an epistatic fitness interaction, where individual loci in one ancestral strain are genetically incompatible with alleles from another strain. We considered other alternatives explanations, some of which were explored in prior work(41). One alternative would be population structure, where recombination may not have had sufficient time to randomly assort alleles after a bottleneck event, though we think that this is an unlikely explanation for our results. One of the key benefits of working with synthetic mapping populations such as RI panels is that the crossing scheme (e.g. fennel cross, round-robin)is designed to minimize such effects(40). Another possibility would be assortative mating, where mate selection drives the selection for or depletion of a particular allele. While possible, and not explicitly arising from epistasis per se, it would still produce a fitness differential across allelic combinations and be picked up by our analysis. Finally, epistatic interactions, the focus of this manuscript, can lead to changes in fitness as demonstrated by prior work(41) which examined Drosophila RI panels in which all expected genotypic combinations for a diallelic cross were generated. They demonstrated a strong correlation between the degree of allelic imbalance and the effect size of the fitness effect of each allelic combination. It follows that similar scenarios should exist in mouse and yeast panels as well. One possible extension of this approach would be to study fitness in the Collaborative Cross strains, another animal population in which a large proportion of the strains went extinct, presumably due to genetic incompatibilities.

In the future, our approach could be leveraged to study additional extant populations or newly created cohorts. For example, our tool could be used to examine whether genetic modifications such as knockouts have differential effects across the mice of a cross, and we are actively exploring its utility in this regard.

## Conclusion

The results of our study suggest that heterogenous SNP effects due to differing ancestries is pervasive in model organism populations. This observation is consistent with prior observations of epistatic interactions in inbred strains of mice as well as examples of ancestry interaction in human studies(10). However, the number and magnitude of ancestry interactions we found was much larger than those found in human studies; we hypothesize this is driven by the larger genetic distance of the ancestors. Further analyses of the sites of polygenic epistasis may reveal novel epistatic interactions which drive phenotypic expression, and suggests that careful attention to genetic ancestry should be considered when studying the role of an individual polymorphism on a phenotype.

## Methods

### Model Organism Populations and Ancestry Calling

Model organism data were drawn from previously reported studies(19,20,30). The LGxSM AIL consists of 1,063 G50-56 mice derived from an original F1 intercross between the LG and SM inbred lines. The Hybrid Mouse Diversity Panel consists of over 150 strains of commercially available inbred mice(27), of which 122 strains were recombinant inbred lines and suitable for our study. The yeast crosses originated from 15 original strains which were bred in pairs to create 15 new crosses. SNPwise ancestries were determined by identifying all SNPs which differed between parental lines (AIL: LG and SM or HMDP: C57BL/6 and A, C3H, DBA/2 or BALB/c or Yeast: See Table S5). Genotypes from the G50-56.RI lines or yeast crosses were filtered for these SNPs and ancestries calculated using either LG or C57BL/6 or one of the yeast strains as the strain background of interest.

### Simulation Framework

#### Power Calculations

We created sets of simulated phenotypes based on the genotypes of the HDMP RI panel, which is an admixed population in which the B6 strain, on average, contributes 50% of each strain’s DNA. For each simulated phenotypes, we drew a SNP (75% > MAF > 25%) at random from the HMDP genotypes and created a phenotype based on β, the genetic effect size, □ the effect size of the interaction between global ancestry (θ) and the chosen SNP and a multivariate normal (mvn) derived from three variance terms: 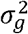, the proportion of variance attributable to genetic effects 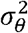, the proportion of variance attributable to Gxθ effects and 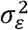, the residual proportional variance attributable to all combined sources of error and variation not considered in this study. Phenotypes were generated both with and without the Gxθ variance term to ascertain the necessity of incorporating a second GRM (**K**^A^), which would correct for relatedness in ancestry, into the algorithm.

We simulated four distinct phenotypes for our analysis

1. Phenotypes generated by including a SNP Effect

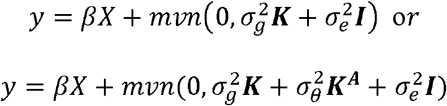
2. Phenotypes generated by including a Gxθ Effect

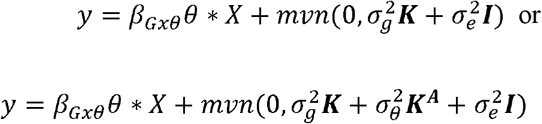

In each phenotype, 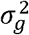 was set to 0.4. When incorporated, 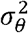 was set to 0.2 and 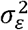 was set to the remainder of the variance (0.6 or 0.4). The power of our model and independence of our *β* and *β_Gxθ_* terms were queried by varying either β or *β_Gxθ_* from 0 to 1 (200 values set 0.005 apart) with 1,000 simulated phenotypes at each step (200,000 total simulations per phenotype).

#### Epistasis Simulation

Using the same panel and set of SNPs described above, we drew one SNP at random to be our test SNP and one to ten additional SNPs to be simulated epistatically interacting SNPs (10,000 simulations per set of interacting SNPs, 100,000 simulations in total). For each simulation we created a composite SNP in which only strains with the minor (non-B6) allele in every one of the tests and interacting SNPs had that allele in the composite SNP. We used this composite SNP to generate a phenotype as described above and selecting an effect size to generate a χ^2^ test statistic of 20.

### Gxθ Model

The equation to determine the effects of a SNP and a SNP x Ancestry term on a phenotype can be written as:

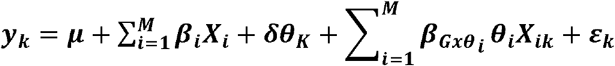

Where *y_k_* is the phenotype of individual k, μ is the mean phenotypic value, M is the number of markers, β are the weights on the SNPs, X is the *m* by *n* array of SNP genotypes, δ is the global weight of the ancestry effect, θ is the ancestries for all N individuals. *β_Gxθ_* are the weights of the Gxθ effect and ε is the combined error term. We want to identify SNPs where *β_Gxθi_* ≠ θ as these are sites where Ancestry is interacting with our genotypes.

Motivated by our model above, we can write a new model for the effect of a single SNP i on a phenotypic trait as:

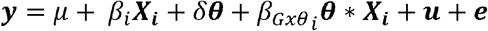

For an individual SNP i. Here, θ is the column vector of ancestries, and θ * X is the element-wise product. The random effect u accounts for relatedness of individuals based on SNPs. Our Gxθ test is then an LRT test with a null of *β_Gxθ_i__* = 0 and an alternate of *β_Gxθ_i__* ≠ 0.

### Reporting Gxθ Results and Identifying Genes of Interest

GWAS thresholds were previously determined to be 4.1E-6 in the case of the HMDP(24) or 8.06E-6 in the case of the AIL(19). An FDR of 5% was used in the yeast crosses as previously reported(25) Genes of Interest were identified by examining nearby genes (within 2 MB) for cis-eQTLs as previously reported in the HMDP(30) and/or by literature and database analyses using bioGPS (www.biogps.org) which draws data from PubMed and other publicly available databases to help researchers understand the role of their genes of interest.

### Determining Significance of Allelic Imbalance Sites

Allelic imbalance significance was determined by performing a Wald test for ancestry frequency (θ) at each SNP i where 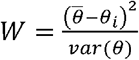 and only strains where the SNP was not invariant were included. Allelic imbalances were considered to be significant after Benjamini-Hochberg correction at a cutoff of 5%.

### Gene Set Enrichment Analysis

All gene enrichments were performed using the GeneAnalytics platform(51), which uses a binomial distribution to assess the enrichment for a user-supplied list of genes within GO terms, Superpaths and other biologically relevant categories. Reported P-values have been corrected for multiple comparisons using the Benjamini-Hochberg correction.

## Declarations

### Ethics Approval and Consent to Participate

Not Applicable

### Consent for Publication

Not Applicable

### Availability of Data and Materials

Data from the HMDP may be accessed at https://systems.genetics.ucla.edu/

Data from the AIL cross may be accessed at http://palmerlab.org/protocols-data/

Data from the Yeast crosses may be accessed at https://github.com/ioshsbloom/veast-16-parents

The Gxθ algorithm may be found at https://github.com/ChristophRau/GxTheta

### Author Contributions

CDR designed the project, created the Gxθ software, acquired the HMDP data and analyzed and interpreted the data used in this manuscript as well as drafted the manuscript. NMG assisted in the design of the AIL cohort, acquired the AIL data and substantially edited the manuscript. JB designed and acquired the yeast cross data and substantially edited the manuscript. DP assisted in conceptualizing the Gxθ algorithm and substantially revised the manuscript. JA provided conceptual support and substantially revised the manuscript. AAP assisted in the design of the AIL cohort and substantially revised the manuscript. AJL assisted in the design of the HMDP cohort and substantially revised the manuscript. NZ conceptualized and designed the Gxθ algorithm, interpreted the data and helped to draft and substantially revise the manuscript. All authors read and approved the final manuscript.

## Acknowledgements

The authors would like to thank Yibin Wang for his constructive critiques of the algorithm and the manuscript.

## Supporting Information Legends

**Figure S1.** Genomic Inflation Factors of SNP and Gxθ associations with increasingly complex SNP-SNP interactions.

**Table S1.** All observed Gxθ associations between clinical traits and SNPs in the AIL, HMDP and Yeast cohorts

**Table S2.** Strains contained in the recombinant inbred panels of the HMDP and their presence in the control or isoproterenol-treated cohorts of the data.

**Table S3** Alleles with significantly altered ancestries at FDR 5% in the HMDP

**Table S4**. Number of SNPs between strains in the AIL and HMDP founder strains as determined by the mouse diversity array. Homozygous differences are 1, heterozygous differences are 0.5

**Table S5.** Information on the yeast crosses used in this manuscript

**Table S6-S9.** Summary statistics for Gxθ results from figure 3

**Table S10.** All significant Gxθ associations for HMDP transcriptome data

