## Supplemental Information for "Modeling epistasis in mice and yeast using the proportion of two or more distinct genetic backgrounds: evidence for “polygenic epistasis”"

**Figure S1.** Lambdas at progressively more complex epistatic interactions for the main and interaction effects as recovered by the GxƟ algorithm

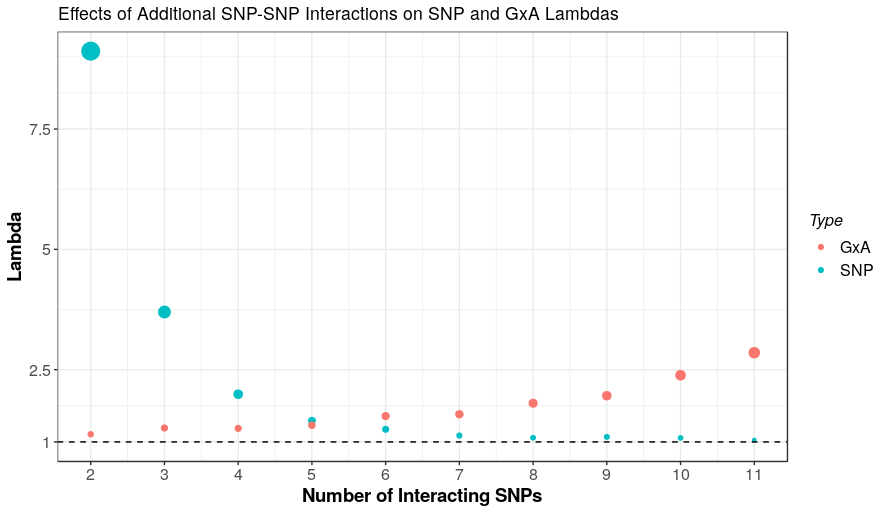

**Table S1.** All observed GxƟ associations between clinical traits and SNPs in the Yeast, AIL, and HMDP cohorts

| **Cohort** | **Phenotype** | **Chr** | **Bp** | **rsID** | **P** |
| --- | --- | --- | --- | --- | --- |
| AIL | act3.2 | 11 | 93607101 | rs27076868 | 6.22E-07 |
| AIL | avg.weight.narm | 6 | 133867165 | rs30057768 | 1.62E-06 |
| AIL | Glucose | 9 | 70318603 | rs50320206 | 3.03E-07 |
| AIL | ppi.weight | 6 | 133899245 | rs47230944 | 2.09E-06 |
| HMDP | E day 21 | 19 | 21360156 | rs36743940 | 5.71E-10 |
| HMDP | E day 21 | 18 | 36333813 | rs31799112 | 1.10E-09 |
| HMDP | E day 21 | 11 | 118264986 | rs26999505 | 1.26E-09 |
| HMDP | E day 21 | 7 | 38211691 | rs32631043 | 1.39E-09 |
| HMDP | E day 21 | 7 | 47678061 | rs49282654 | 1.84E-09 |
| HMDP | E day 21 | 4 | 144572533 | rs27616794 | 2.43E-09 |
| HMDP | E day 21 | 3 | 26466861 | rs38357506 | 2.44E-09 |
| HMDP | E day 21 | 1 | 63943836 | rs30079554 | 4.84E-08 |
| HMDP | E day 21 | 1 | 111320857 | rs48448024 | 1.29E-07 |
| HMDP | E day 21 | 2 | 70920690 | rs27971838 | 1.23E-08 |
| HMDP | E day 21 | 3 | 129615106 | rs16790998 | 2.52E-08 |
| HMDP | E day 21 | 4 | 134970895 | rs27619494 | 1.42E-08 |
| HMDP | E day 21 | 5 | 35419668 | rs32537851 | 7.45E-08 |
| HMDP | E day 21 | 5 | 120017310 | rs52255477 | 7.63E-08 |
| HMDP | E day 21 | 10 | 88298448 | rs45869923 | 4.06E-08 |
| HMDP | E day 21 | 11 | 7182347 | rs26897032 | 3.99E-08 |
| HMDP | E day 21 | 12 | 114144765 | rs47800412 | 2.64E-08 |
| HMDP | E day 21 | 13 | 72333688 | rs46580133 | 5.45E-08 |
| HMDP | E day 21 | 15 | 95123948 | rs46936246 | 1.97E-08 |
| HMDP | E day 21 | 14 | 10864544 | rs30617637 | 2.69E-08 |
| HMDP | E day 21 | 17 | 29523029 | rs33612883 | 8.75E-08 |
| HMDP | E day 21 | X | 129292043 | rs29272711 | 1.22E-08 |
| HMDP | E/A Ratio Day 21 | 6 | 8462274 | rs47261338 | 7.10E-08 |
| HMDP | E/A Ratio Day 21 | 13 | 10173630 | rs48793742 | 7.02E-07 |
| HMDP | FFA Control | 11 | 113035483 | rs27020574 | 4.09E-06 |
| HMDP | Fibrosis Control | 17 | 41256115 | rs6295287 | 6.74E-07 |
| HMDP | Fibrosis Control | 19 | 61215984 | rs30715237 | 1.55E-06 |
| HMDP | Fibrosis Control | 14 | 18292706 | rs31538570 | 2.14E-06 |
| HMDP | Fibrosis Control | 7 | 134530651 | rs33146511 | 2.54E-06 |
| HMDP | FS Day 0 | X | 48286693 | rs30272504 | 3.69E-06 |
| HMDP | Heart Rate Day 0 | X | 48286693 | rs30272504 | 2.68E-06 |
| HMDP | Heart Rate Day 7 | 18 | 9408717 | rs29769121 | 2.37E-06 |
| HMDP | Heart Rate Day 7 | 5 | 98753288 | rs31910795 | 2.60E-06 |
| HMDP | LVIDs Day 0 | X | 48286693 | rs30272504 | 2.92E-06 |
| HMDP | LVM Day 14 | 17 | 72564961 | rs50549031 | 8.65E-08 |
| HMDP | LVM Day 14 | 7 | 66027128 | rs33004129 | 1.10E-07 |
| HMDP | LVM Day 14 | X | 10000011 | rs33501756 | 1.17E-06 |
| HMDP | LVM Day 7 | 3 | 148140470 | rs31313229 | 2.91E-06 |
| HMDP | LVMc Day 7 | 3 | 148140470 | rs31313229 | 2.91E-06 |
| HMDP | wLA ISO | 16 | 54550810 | rs51319671 | 2.18E-06 |
| HMDP | wLA ISO | 4 | 46325541 | rs27816179 | 2.46E-06 |
| HMDP | wRA ISO | 16 | 49837267 | rs50479702 | 4.13E-07 |
| HMDP | wRA ISO | 6 | 73857753 | rs38199107 | 1.95E-06 |
| HMDP | wRA ISO | 4 | 46802557 | rs27800512 | 2.49E-06 |
| Yeast | Cross 375: Fluconazole | II | 309402 | NA | 0.000119 |
| Yeast | Cross 375: Fluconazole | III | 266399 | NA | 0.000145 |
| Yeast | Cross 375: Fluconazole | VI | 129958 | NA | 6.54E-06 |
| Yeast | Cross 375: Fluconazole | XI | 641793 | NA | 0.000224 |
| Yeast | Cross 375: Methotrexate | II | 260 | NA | 7.92E-05 |
| Yeast | Cross 375: Methotrexate | VII | 1005304 | NA | 2.15E-05 |
| Yeast | Cross 377: Maltose | VII | 1067384 | NA | 7.98E-05 |
| Yeast | Cross 377: Xylose | X | 272177 | NA | 7.67E-06 |
| Yeast | Cross 381: Glycerol | XII | 573838 | NA | 3.97E-06 |
| Yeast | Cross 381: YNB | IV | 1170023 | NA | 3.69E-06 |
| Yeast | Cross 381: YNB | XIII | 275756 | NA | 0.000364 |
| Yeast | Cross 381: Zeocin | VIII | 20921 | NA | 3.65E-06 |
| Yeast | Cross 393: Copper_Sulfate | XII | 578550 | NA | 3.38E-06 |
| Yeast | Cross 393: Tunicamycin | V | 246808 | NA | 7.52E-06 |
| Yeast | Cross 2999: EGTA | V | 219603 | NA | 6.51E-05 |
| Yeast | Cross 2999: EGTA | XIV | 622639 | NA | 2.91E-07 |
| Yeast | Cross 2999: Lithium_Chloride | II | 105865 | NA | 0.001615 |
| Yeast | Cross 2999: Lithium_Chloride | IV | 519675 | NA | 1.04E-07 |
| Yeast | Cross 2999: Lithium_Chloride | IX | 38806 | NA | 1.31E-05 |
| Yeast | Cross 2999: Lithium_Chloride | VI | 256157 | NA | 0.000629 |
| Yeast | Cross 2999: Lithium_Chloride | VII | 62447 | NA | 9.92E-05 |
| Yeast | Cross 2999: Lithium_Chloride | VIII | 359999 | NA | 0.001896 |
| Yeast | Cross 2999: Lithium_Chloride | X | 97232 | NA | 6.05E-05 |
| Yeast | Cross 2999: Lithium_Chloride | XII | 341678 | NA | 1.16E-05 |
| Yeast | Cross 2999: Lithium_Chloride | XVI | 447414 | NA | 0.000114 |
| Yeast | Cross 2999: Manganese_Sulfate | XIV | 622639 | NA | 2.30E-06 |
| Yeast | Cross 3000: 6-azauracil | IV | 1384392 | NA | 5.75E-05 |
| Yeast | Cross 3000: 6-azauracil | XIII | 159493 | NA | 1.80E-05 |
| Yeast | Cross 3000: EtOH_Glucose | IV | 1381584 | NA | 5.50E-05 |
| Yeast | Cross 3000: EtOH_Glucose | XIII | 131108 | NA | 1.51E-06 |
| Yeast | Cross 3001: Formamide | X | 517094 | NA | 9.81E-07 |
| Yeast | Cross 3003: Copper_Sulfate | XV | 83697 | NA | 2.72E-06 |
| Yeast | Cross 3003: EGTA | IX | 53940 | NA | 4.52E-05 |
| Yeast | Cross 3003: EGTA | XII | 410115 | NA | 5.76E-05 |
| Yeast | Cross 3003: SDS | VIII | 63044 | NA | 5.35E-06 |
| Yeast | Cross 3003: SDS | X | 59783 | NA | 0.001324 |
| Yeast | Cross 3003: SDS | XVI | 311427 | NA | 4.73E-07 |
| Yeast | Cross 3003: YNB | XV | 251169 | NA | 7.81E-07 |
| Yeast | Cross 3008: Copper_Sulfate | VIII | 216235 | NA | 2.77E-06 |
| Yeast | Cross 3008: Magnesium_Chloride | VIII | 62702 | NA | 1.22E-05 |
| Yeast | Cross 3008: Magnesium_Chloride | XVI | 220547 | NA | 1.25E-05 |
| Yeast | Cross 3028: Cadmium_Chloride | II | 794828 | NA | 8.33E-06 |
| Yeast | Cross 3028: Cadmium_Chloride | XVI | 169035 | NA | 6.50E-07 |
| Yeast | Cross 3028: EtOH_Glucose | IV | 1387173 | NA | 4.56E-05 |
| Yeast | Cross 3028: EtOH_Glucose | XI | 143378 | NA | 2.19E-06 |
| Yeast | Cross 3028: EtOH_Glucose | XIII | 858543 | NA | 4.79E-05 |
| Yeast | Cross 3028: Fructose | IV | 241065 | NA | 2.39E-05 |
| Yeast | Cross 3028: Fructose | XI | 143378 | NA | 6.76E-05 |
| Yeast | Cross 3028: Fructose | XII | 418184 | NA | 4.65E-05 |
| Yeast | Cross 3028: Fructose | XV | 292642 | NA | 0.000151 |
| Yeast | Cross 3028: Galactose | IV | 241065 | NA | 1.47E-05 |
| Yeast | Cross 3028: Galactose | V | 402722 | NA | 2.01E-05 |
| Yeast | Cross 3028: Galactose | XII | 418184 | NA | 3.87E-05 |
| Yeast | Cross 3028: Mannose | IV | 241065 | NA | 7.15E-05 |
| Yeast | Cross 3028: Mannose | VI | 96304 | NA | 0.000134 |
| Yeast | Cross 3028: Mannose | XI | 143378 | NA | 2.80E-05 |
| Yeast | Cross 3028: Mannose | XII | 418844 | NA | 1.97E-05 |
| Yeast | Cross 3028: Mannose | XV | 443031 | NA | 0.000215 |
| Yeast | Cross 3028: Mannose | XVI | 334546 | NA | 0.000193 |
| Yeast | Cross 3028: Trehalose | IX | 164730 | NA | 4.48E-05 |
| Yeast | Cross 3028: Trehalose | XI | 158228 | NA | 7.68E-06 |
| Yeast | Cross 3028: Xylose | XI | 141291 | NA | 9.55E-06 |
| Yeast | Cross 3028: YPD | IV | 1387173 | NA | 7.26E-06 |
| Yeast | Cross 3043: Galactose | II | 238655 | NA | 2.80E-05 |
| Yeast | Cross 3043: Galactose | XI | 147108 | NA | 3.30E-05 |
| Yeast | Cross 3043: Galactose | XV | 561457 | NA | 0.000498 |
| Yeast | Cross 3043: Manganese_Sulfate | XIII | 539307 | NA | 9.60E-05 |
| Yeast | Cross 3043: Manganese_Sulfate | XVI | 206053 | NA | 4.17E-10 |
| Yeast | Cross 3043: Paraquat | XV | 410166 | NA | 3.14E-08 |
| Yeast | Cross 3043: YNB | XV | 561457 | NA | 3.67E-07 |
| Yeast | Cross 3043: YPD | XV | 561457 | NA | 2.50E-06 |
| Yeast | Cross 3043: Zeocin | V | 10492 | NA | 0.00042 |
| Yeast | Cross 3043: Zeocin | VII | 1063697 | NA | 0.00049 |
| Yeast | Cross 3043: Zeocin | VIII | 17612 | NA | 7.15E-05 |
| Yeast | Cross 3043: Zeocin | X | 571909 | NA | 1.56E-05 |
| Yeast | Cross 3049: Paraquat | I | 90039 | NA | 0.000587 |
| Yeast | Cross 3049: Paraquat | II | 636937 | NA | 0.004824 |
| Yeast | Cross 3049: Paraquat | III | 213865 | NA | 0.002355 |
| Yeast | Cross 3049: Paraquat | IV | 227289 | NA | 8.81E-05 |
| Yeast | Cross 3049: Paraquat | IX | 118855 | NA | 0.004751 |
| Yeast | Cross 3049: Paraquat | VII | 705208 | NA | 0.001372 |
| Yeast | Cross 3049: Paraquat | VIII | 77756 | NA | 8.55E-06 |
| Yeast | Cross 3049: Paraquat | X | 28329 | NA | 3.20E-05 |
| Yeast | Cross 3049: Paraquat | XI | 115636 | NA | 6.33E-05 |
| Yeast | Cross 3049: Paraquat | XII | 552497 | NA | 2.61E-05 |
| Yeast | Cross 3049: Paraquat | XIII | 510798 | NA | 0.000313 |
| Yeast | Cross 3049: Paraquat | XV | 185029 | NA | 0.001455 |
| Yeast | Cross 3049: Paraquat | XVI | 581602 | NA | 0.000421 |
| Yeast | Cross A: SDS | XIII | 84233 | NA | 1.73E-05 |
| Yeast | Cross B: Maltose | VII | 1067754 | NA | 1.20E-07 |
| Yeast | Cross B: Maltose | XII | 331594 | NA | 6.40E-05 |
| Yeast | Cross B: Sorbitol | VII | 1068934 | NA | 3.78E-05 |

**Table S2.** Strains contained in the recombinant inbred panels of the HMDP and their presence in the control or isoproterenol-treated cohorts of the data.

| **Panel** | **Strain** | **Control** | **Treated** |
| --- | --- | --- | --- |
| AxB | AXB1/PgnJ |  |  |
| AxB | AXB10/PgnJ | X | X |
| AxB | AXB12/PgnJ | X |  |
| AxB | AXB13/PgnJ |  |  |
| AxB | AXB15/PgnJ |  |  |
| AxB | AXB19/PgnJ | X | X |
| AxB | AXB19a/PgnJ | X | X |
| AxB | AXB19b/PgnJ | X | X |
| AxB | AXB2/PgnJ |  |  |
| AxB | AXB23/PgnJ |  |  |
| AxB | AXB24/PgnJ |  |  |
| AxB | AXB4/PgnJ | X | X |
| AxB | AXB5/PgnJ |  |  |
| AxB | AXB6/PgnJ | X | X |
| AxB | AXB8/PgnJ | X | X |
| AxB | BXA1/PgnJ | X | X |
| AxB | BXA11/PgnJ | X | X |
| AxB | BXA12/PgnJ | X |  |
| AxB | BXA13/PgnJ |  |  |
| AxB | BXA14/PgnJ | X | X |
| AxB | BXA16/PgnJ | X | X |
| AxB | BXA2/PgnJ | X | X |
| AxB | BXA24/PgnJ | X | X |
| AxB | BXA25/PgnJ |  |  |
| AxB | BXA26/PgnJ |  |  |
| AxB | BXA4/PgnJ | X | X |
| AxB | BXA7/PgnJ | X | X |
| AxB | BXA8/PgnJ | X | X |
| BxD | BXD1/TyJ | X |  |
| BxD | BXD100/RwwJ |  |  |
| BxD | BXD102/RwwJ |  |  |
| BxD | BXD11/TyJ | X | X |
| BxD | BXD12/TyJ | X | X |
| BxD | BXD13/TyJ |  |  |
| BxD | BXD14/TyJ | X | X |
| BxD | BXD15/TyJ | X |  |
| BxD | BXD16/TyJ |  |  |
| BxD | BXD18/TyJ |  |  |
| BxD | BXD19/TyJ |  | X |
| BxD | BXD2/TyJ |  |  |
| BxD | BXD20/TyJ |  |  |
| BxD | BXD21/TyJ | X | X |
| BxD | BXD22/TyJ | X | X |
| BxD | BXD24b/TyJ |  |  |
| BxD | BXD27/TyJ | X |  |
| BxD | BXD28/TyJ |  |  |
| BxD | BXD29/TyJ |  |  |
| BxD | BXD31/TyJ | X | X |
| BxD | BXD32/TyJ | X | X |
| BxD | BXD33/TyJ | X |  |
| BxD | BXD34/TyJ | X |  |
| BxD | BXD36/TyJ |  |  |
| BxD | BXD38/TyJ | X | X |
| BxD | BXD39/TyJ | X | X |
| BxD | BXD40/TyJ | X | X |
| BxD | BXD42/TyJ |  |  |
| BxD | BXD43/RwwJ | X | X |
| BxD | BXD44/RwwJ | X | X |
| BxD | BXD45/RwwJ | X | X |
| BxD | BXD48/RwwJ | X | X |
| BxD | BXD49/RwwJ | X | X |
| BxD | BXD5/TyJ | X | X |
| BxD | BXD50/RwwJ | X | X |
| BxD | BXD51/RwwJ |  |  |
| BxD | BXD55/RwwJ | X | X |
| BxD | BXD56/RwwJ | X | X |
| BxD | BXD6/TyJ | X | X |
| BxD | BXD60/RwwJ |  |  |
| BxD | BXD61/RwwJ | X | X |
| BxD | BXD63/RwwJ | X | X |
| BxD | BXD64/RwwJ | X | X |
| BxD | BXD65/RwwJ |  |  |
| BxD | BXD66/RwwJ | X | X |
| BxD | BXD67/RwwJ |  |  |
| BxD | BXD68/RwwJ | X | X |
| BxD | BXD69/RwwJ | X |  |
| BxD | BXD70/RwwJ | X | X |
| BxD | BXD71/RwwJ | X | X |
| BxD | BXD73/RwwJ | X | X |
| BxD | BXD74/RwwJ | X | X |
| BxD | BXD75/RwwJ | X | X |
| BxD | BXD77/RwwJ |  |  |
| BxD | BXD79/RwwJ | X | X |
| BxD | BXD8/TyJ | X | X |
| BxD | BXD80/RwwJ |  |  |
| BxD | BXD81/RwwJ |  |  |
| BxD | BXD84/RwwJ | X | X |
| BxD | BXD85/RwwJ | X | X |
| BxD | BXD86/RwwJ | X |  |
| BxD | BXD87/RwwJ | X | X |
| BxD | BXD89/RwwJ |  |  |
| BxD | BXD9/TyJ |  |  |
| BxD | BXD90/RwwJ |  |  |
| BxD | BXD92/RwwJ |  |  |
| BxD | BXD95/RwwJ |  |  |
| BxD | BXD96/RwwJ |  |  |
| BxD | BXD97/RwwJ |  |  |
| BxD | BXD98/RwwJ |  |  |
| BxD | BXD99/RwwJ |  |  |
| BxH | BXH10/TyJ |  |  |
| BxH | BXH11/TyJ |  |  |
| BxH | BXH14/TyJ |  |  |
| BxH | BXH19/TyJ | X | X |
| BxH | BXH2/TyJ |  |  |
| BxH | BXH20/KccJ |  |  |
| BxH | BXH22/KccJ |  |  |
| BxH | BXH4/TyJ |  | X |
| BxH | BXH6/TyJ | X | X |
| BxH | BXH7/TyJ |  |  |
| BxH | BXH8/TyJ | X | X |
| BxH | BXH9/TyJ | X | X |
| CxB | CXB1/ByJ |  |  |
| CxB | CXB11/HiAJ | X | X |
| CxB | CXB12/HiAJ | X | X |
| CxB | CXB13/HiAJ | X | X |
| CxB | CXB2/ByJ |  |  |
| CxB | CXB3/ByJ | X | X |
| CxB | CXB4/ByJ |  |  |
| CxB | CXB6/ByJ | X | X |
| CxB | CXB7/ByJ | X | X |
| CxB | CXB8/HiAJ | X | X |
| CxB | CXB9/HiAJ |  |  |

**Table S3** Alleles with significantly altered ancestries at FDR 5% in the HMDP

**Table S4**. Number of SNPs between strains in the AIL and HMDP founder strains as determined by the mouse diversity array. Homozygous differences are 1, heterozygous differences are 0.5

| **Strain 1** | **Strain 2** | **Differences** |
| --- | --- | --- |
| A/J | C57BL/6J | 112872.5 |
| C57BL/6J | DBA/2J | 112536.5 |
| C3H/HeJ | C57BL/6J | 112052.5 |
| C57BL/6J | SM/J | 108583.5 |
| C57BL/6J | LG/J | 107193 |
| DBA/2J | LG/J | 102927.5 |
| BALB/cJ | C57BL/6J | 101991.5 |
| A/J | SM/J | 99919 |
| A/J | DBA/2J | 99457.5 |
| LG/J | SM/J | 98463 |
| DBA/2J | SM/J | 97388.5 |
| BALB/cJ | SM/J | 96681.5 |
| BALB/cJ | DBA/2J | 96352.5 |
| C3H/HeJ | SM/J | 94627 |
| C3H/HeJ | LG/J | 89901.5 |
| A/J | LG/J | 85687.5 |
| BALB/cJ | LG/J | 83197 |
| C3H/HeJ | DBA/2J | 79223.5 |
| BALB/cJ | C3H/HeJ | 74288 |
| A/J | C3H/HeJ | 65204 |
| A/J | BALB/cJ | 56490.5 |

**Table S5**. Information on the Yeast Crosses used in this manuscript

| **Cross** | **Parent 1** | **Parent 2** |
| --- | --- | --- |
| 375 | BYa | M22 |
| A | ByA | RMx |
| 376 | RMx | YPS163a |
| B | YJM145x | YPS163a |
| 377 | CLIB413a | YJM145x |
| 393 | CLIB413a | YJM978x |
| 381 | YJM454a | YJM978x |
| 3008 | YJM454a | YPS1009x |
| 2999 | I14a | YPS1009x |
| 3000 | I14a | Y10x |
| 3001 | PW5a | Y10x |
| 3049 | 273614xa | PW5a |
| 3003 | 273614xa | YJM981x |
| 3043 | CBS2888a | CLIB219x |
| 3028 | CLIB219x | M22 |

**Table S6.** Summary statistics for GxƟ analyses in figure 3 part A

**Table S7.** Summary statistics for GxƟ analyses in figure 3 part B.

**Table S8.** Summary statistics for GxƟ analyses in figure 3 part C.

**Table S9.** Summary statistics for GxƟ analyses in figure 3 part D

**Table S10.** Bonferroni-corrected genome-wide significant GxƟ associations between gene expression in control and isoproterenol-treated mice of the HMDP
